# A reference brain for the clonal raider ant

**DOI:** 10.1101/2025.10.13.679875

**Authors:** Dominic D. Frank, Lindsey E. Lopes, Rishika Mohanta, Isabelle Seckler, Ivan Lacroix, Daniel J. C. Kronauer

## Abstract

Ants exhibit remarkable collective and social behaviors, such as alloparental care^1^, chemical communication^2^, homing^3^, and cooperative group hygiene^4^. The clonal raider ant *Ooceraea biroi* is especially well-suited for investigating the neuronal and genetic underpinnings of these behaviors^5^. Unlike most ant species, *O. biroi* lacks a queen caste. Instead, colonies consist entirely of regular workers and slightly larger intercaste workers^6^. All workers reproduce in synchrony via parthenogenesis, giving rise to age-matched cohorts of clonally identical offspring^7,8^. This unique life history enables precise experimental control over age, genotype, and colony composition. These features have also facilitated the introduction of genetically encoded calcium indicators into *O. biroi*, enabling *in vivo* two-photon imaging to investigate the neural basis of social behaviors^9^. Despite its promise as a neuroscience model, the structure of the clonal raider ant brain has not been systematically characterized, and a representative reference brain does not exist. To address this gap, we imaged the brains of 40 age-matched, genetically identical individuals with confocal microscopy and, using 3D groupwise registration, generated the first reference brain for the species. We introduce a registration pipeline to align brains to this reference, facilitating the comparison of anatomical features across labeling experiments with high spatial precision. Unexpectedly, despite homogeneity in genotype, age, and external morphology, we discovered extensive interindividual variability across our collection of brain samples. This raises the possibility that behavioral division of labor in *O. biroi* is linked to individual differences in brain structure. This work provides a powerful resource for the emerging clonal raider ant neuroscience community and reveals novel features of the species’ neurobiology that may influence social behaviors and colony function.

## RESULTS

### A clonal raider ant reference brain reveals asymmetries in brain anatomy

Reference brains are valuable tools for studying the central nervous system. They provide a framework for comparing imaging experiments^10–13^, serve as atlases^14^, act as anatomical scaffolds for connectomics data^10,11,15,16^, and support online resources, such as the Virtual Fly Brain and Insect Brain Database^17,18^.

To generate the clonal raider ant reference brain, we used 3D groupwise registration similar to that used in *Drosophila melanogaster*^10,19^. This approach aligns multiple brains to produce an unbiased template reflecting population-level anatomy. We labeled the neuropil of 40 one-month-old female *O. biroi* clonal line B (a well-characterized laboratory genotype^7^) workers using a monoclonal antibody that recognizes the presynaptic protein Synapsin and imaged their brains with confocal microscopy at 0.1289 × 0.1289 × 0.295 μm³ voxel resolution (**Figure 1A**).

**Figure 1.**
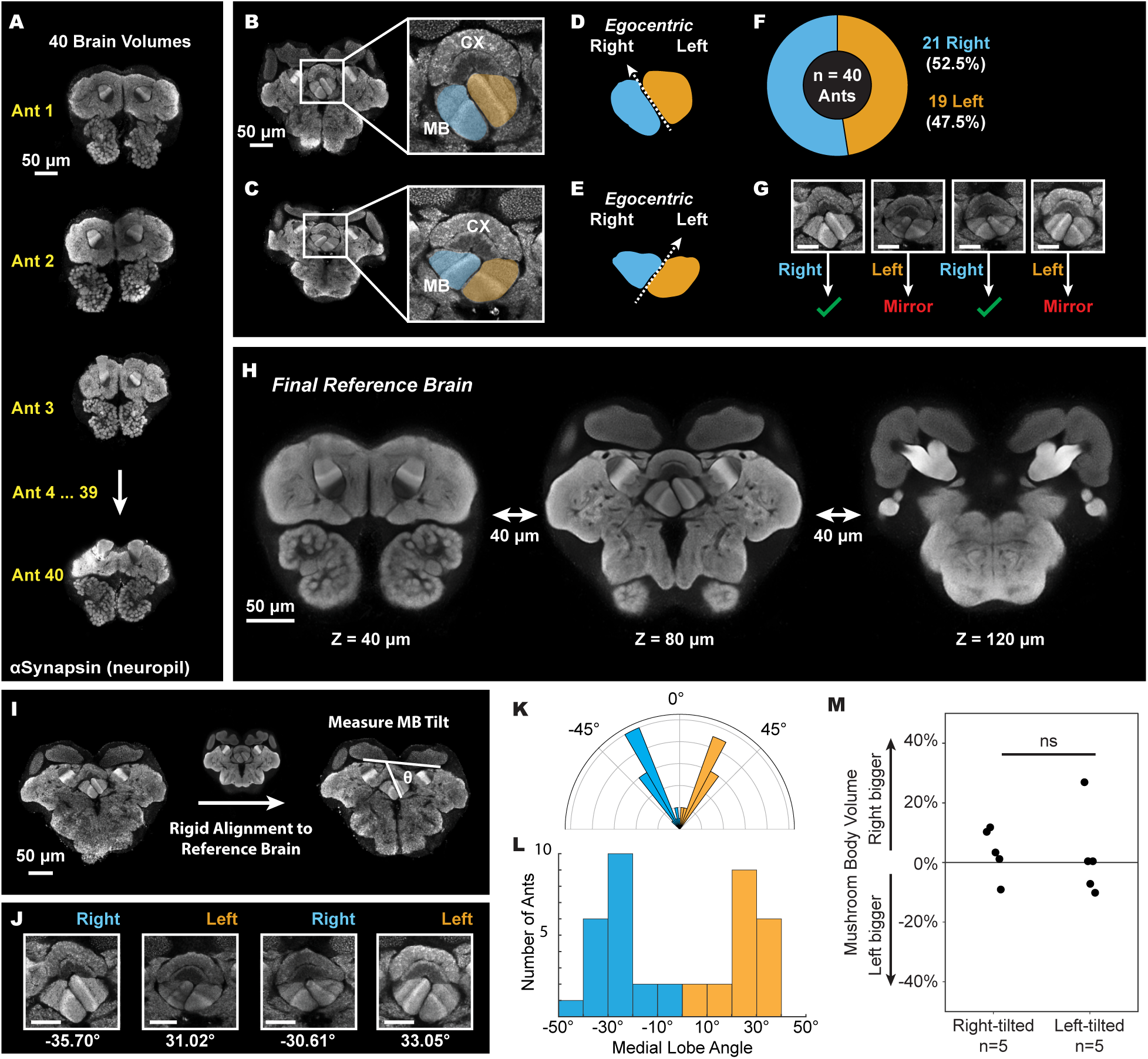
A clonal raider ant reference brain reveals asymmetries in brain anatomy. (A) Example confocal images from the 40 individual clonal raider ant brains used to generate the reference brain. Brain neuropil is labeled with an anti-Synapsin antibody. Image planes are from the same position in each brain. (B-C) Confocal images of two clonal raider ant brains with either a right (B) or left (C) tilt. The right and left mushroom body (MB) medial lobes are highlighted in blue and orange, respectively. Image planes are from a position posterior to those shown in (A). (D-E) Cartoon diagrams demonstrating the ‘right’ and ‘left’ MB tilt. (F) Counts of right- and left-tilted brains among the 40 brains used to generate the reference brain. (G) MB medial lobes from four additional example brains and their classification as right- or left-tilted. To avoid artifacts in the reference brain, all left-tilted brains were mirrored prior to registration. Scale bars are 25μm. (H) Z-slices from the clonal raider ant reference brain, each 40μm apart in the anterior-posterior axis. The left image corresponds to the position in (A), the center image corresponds to the position in (B-C), and the right image shows a more posterior position. See Supplemental Figure 1 for a flowchart detailing reference brain construction. (I) Illustration of the approach for quantifying mushroom body tilt. (J) Confocal images of example mushroom body medial lobes from Figure 1G with the measured angle indicated. Scale bars are 25μm. (K) Polar plot of mushroom body angles measured in the 40 individual clonal raider ant brains used to generate the reference brain. (L) Histogram of the mushroom body angles. Same data as in (K). (M) Relative size of left and right mushroom bodies in left-tilted (n = 5) and right-tilted (n = 5) brains. Each data point represents the right mushroom body volume divided by the left mushroom body volume for a single brain. Medians were compared with the Mann-Whitney U test (p = 0.4206).

While inspecting these brains, we identified an obvious structural asymmetry in the tilt of the mushroom body (MB) medial lobes at the brain midline. The tilt direction varied from individual to individual (**Figure 1B-C**). We categorized each brain as egocentric right-tilted (n=21) or left-tilted (n=19) (**Figure 1D-F**) and mirrored the left-tilted brains to prevent reference artifacts (**Figure 1G**). We next used Advanced Normalization Tools (ANTs^20,21^) to construct the reference using groupwise registration and averaging of the 40 individual brains. Reference generation in ANTs first uses affine registration, which adjusts for differences in size, position, and orientation, followed by iterative rounds of diffeomorphic registration, which enables more flexible, non-linear alignment **(Supplemental Figure 1A)**^22^. Together, these steps yielded a high-resolution reference brain for *O. biroi* (**Figure 1H**). Irregularities present in the individual brains were absent in the final reference brain and neuropil boundaries remained clear and sometimes more pronounced (*e.g.,* compare **Figure 1A** and the left-most image in **Figure 1H**).

To characterize the anatomy of the reference brain, we reconstructed and quantified volumes for 11 focal regions that could be unambiguously identified in our anti-Synapsin channel: the bilateral mushroom bodies (MBs), antennal lobes (ALs), and optic lobes (OLs), plus the central complex (CX), which includes paired noduli (NO), upper and lower central bodies (CBU, CBL), and the protocerebral bridge (PB) (**Supplemental Figure 1B-F**). For comparison, we also quantified volumes for the same neuropils in *D. melanogaster* using the JRC2018 brain template (**Supplemental Figure 1G**). In clonal raider ants, which lack eyes and rely heavily on olfaction, the ALs and MBs occupy ∼13% and ∼25% of total brain volume, respectively, compared to ∼4% and ∼3% in *D. melanogaster*. The optic lobes, on the other hand, are heavily reduced (∼0.5%) compared to fruit flies (∼39%). The proportionate volume of the CX is approximately half that of flies (∼1% vs. ∼2%) (**Supplemental Figure 1G**).

As a first experiment using the reference brain, we sought to quantify the MB medial lobe tilt described above. We aligned each of the 40 individual brains to the reference brain with a rigid registration, which brings the brains into a common orientation without distorting anatomical structures (**Figure 1I**). Next, we identified an approximately equivalent Z-plane in each registered brain and measured the angle of the tilt (**Figure 1J**). We observed a bimodal distribution with the two peaks centered on approximately +/− 30°, confirming our initial observation that the clonal raider ant MB medial lobe displays one of two distinct phenotypes (**Figure 1K-L**). To determine if MB volume varied with tilt direction, we segmented bilateral MBs for a selection of individual brains. The right-to-left MB volume ratio did not differ between right- and left-tilted brains (**Figure 1M**), indicating the two hemispheres vary in orientation but not overall size.

### The brains of clonal raider ants vary in size

Despite controlling for age and genotype, total brain volume varied more than two-fold across the 40 individual brains used to generate the reference (**Figure 2A-B**). Brains with left- and right-tilted MBs had similar volumes (**Supplemental Figure 2A**), implying multiple independent sources of variability. To assess whether volume variability is specific to the clonal line B genetic background, we immunolabeled and imaged brains of one-month-old individuals of another clonal genotype, line A^7^. The variability in brain sizes across these two strains was similar (**Supplemental Figure 2B**), indicating that this phenomenon is not lineage-specific.

**Figure 2.**
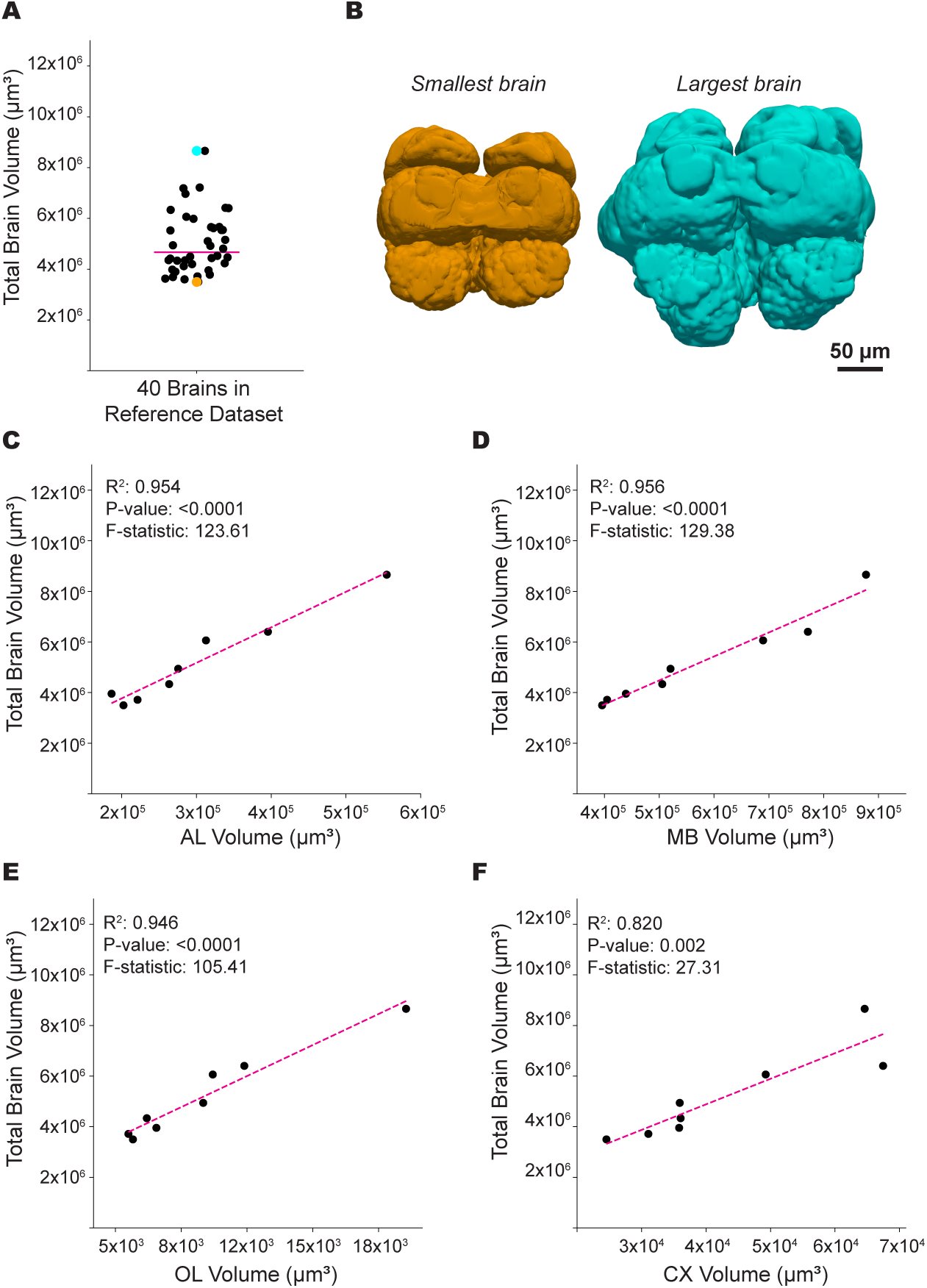
The brains of clonal raider ants vary in size. (A) Total brain volume in cubic micrometers for the 40 brains used to create the reference brain. Each data point represents a single brain; the magenta line indicates the median. Orange and blue data points correspond to the smallest and largest brain in the dataset, respectively. (B) 3D renderings of the smallest and largest brain in our reference brain dataset. Colors correspond to the points in (A). (C-F) Linear regression plots of total brain volume (Y axis) and the volume of focal neuropils (X axis) for 8 individual brains. Focal neuropils include (C) antennal lobes, AL; (D) mushroom bodies, MB; (E) optic lobes, OL; and (F) central complex, CX. Each data point represents a single brain; the magenta line represents the linear model. Neuropil volume is the mean of both hemispheres for bilateral neuropils.

To test whether the variation arises from specific brain regions, as has been observed in *Camponotus* and *Cataglyphis* ants, where certain neuropils expand with age or changes in behavioral role^23,24^, we manually segmented the ALs, MBs, CX, and OLs in a subset of the 40 individual brains used to generate the reference. Each of these neuropils scaled proportionally with total brain volume (**Figure 2C-F**), indicating that the variation in overall brain size is not due to neuropil-specific expansions.

Next, we examined possible body size and caste effects on brain volume by comparing the brains of one-month-old regular workers and intercaste workers from clonal line B. Intercastes are large workers with traits intermediate between regular workers and queens. They have larger ovaries, rudimentary eyespots, and subtle fissures on the mesonotum^6^. Despite these differences, brain volumes and variability were similar in both groups (**Supplemental Figure 2D**). Finally, in 50 additional regular workers, body size and head area did not correlate with brain volume (**Supplemental Figure 2D–F**). These results suggest that brain size is highly variable in clonal raider ants, without being strongly influenced by body size or caste phenotype.

### Registrations to the reference brain are precise

Any brain labeled with the anti-Synapsin antibody can be registered to the reference brain, and the same transformation can be applied to additional channels, enabling 3D visualization in a shared virtual reference space. As a proof of concept, we immunostained brains using a panel of 14 antibodies that label major neurotransmitter systems or cytological features (*e.g.,* axon tracts) in insects, of which four produced specific labeling (see **Table S1** for all antibodies tested and anatomical descriptions). These antibodies recognize gamma-aminobutyric acid (GABA), serotonin (5-HT), tyrosine hydroxylase (TH; a proxy for dopamine), and inotocin, the insect ortholog of oxytocin/vasopressin. We labeled brains with one of these four antibodies alongside the anti-Synapsin antibody and collected two-channel confocal volumes.

Next, we developed a registration pipeline supported by a user-friendly GUI (see Methods). Samples with left-tilted MBs were mirrored to match the reference. The anti-Synapsin “reference” channel was then registered to the reference brain using an affine transformation followed by a diffeomorphic transformation. Finally, the same transformations were applied to the “experimental” channel. **Figure 3A-C** shows registered example brains.

**Figure 3.**
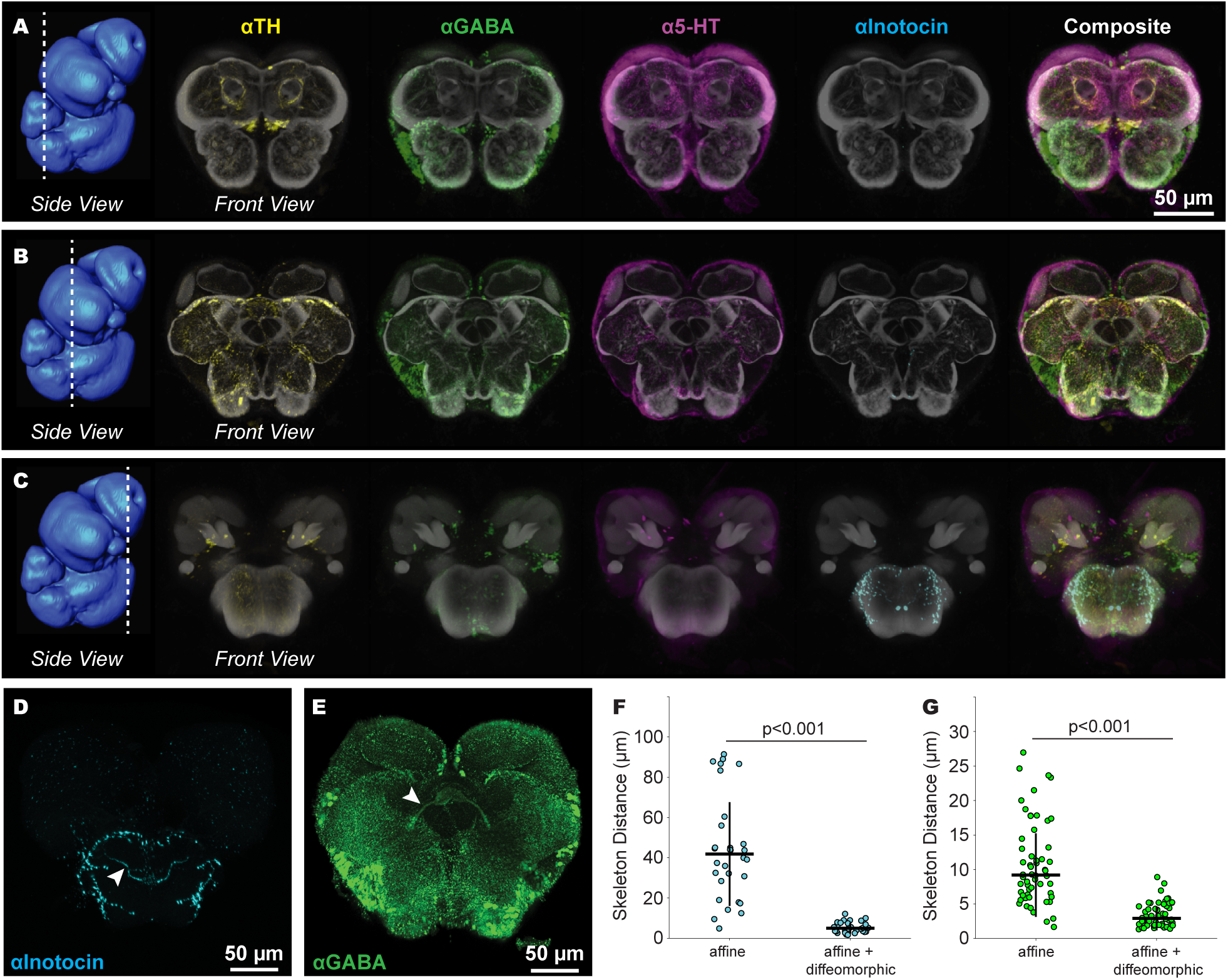
Registrations to the reference brain are precise. (A-C) Four example brains (columns), each immunostained for a different neurotransmitter and registered to the reference brain. Each row contains images from a different Z-plane, indicated by the dashed lines overlaid on the 3D rendering of the reference brain to the left. The composite (right column) shows all four neurotransmitter labels overlaid. The anti-Synapsin channel is displayed in greyscale for reference. (D-E) Representative Z-planes from brains stained with (D) anti-inotocin (n = 6 brains) or (E) anti-GABA antibodies (n = 8 brains) and registered to the reference brain. Arrowheads indicate neurite tracts that were manually skeletonized bilaterally (n = 12 inotocin skeletons; n = 16 GABA skeletons). (F-G) Pairwise mean distances between inotocin (F) and GABA (G) skeletons, calculated after registration using either affine transformations alone (left) or affine plus diffeomorphic transformations (right). All pairwise comparisons were performed between skeletons from the same brain hemisphere (right–right and left–left), giving 30 total comparisons for inotocin (15 per side) and 56 total comparisons for GABA (28 per side). Medians were compared using two-tailed Welch’s t-tests. Plots show medians (horizontal lines) and standard deviations (vertical lines). Abbreviations: tyrosine hydroxylase (TH), gamma-aminobutyric acid (GABA), 5-hydroxytryptamine (5-HT).

To quantify registration precision, we used brains labeled with the anti-inotocin and anti-GABA antibodies. These antibodies mark distinct fiber tracts, allowing for skeletonization and pairwise distance measurements (**Figure 3D–E**). We bilaterally skeletonized these tracts in brains registered using either (1) an affine transformation alone or (2) our standard registration pipeline (*i.e.,* affine transformation followed by diffeomorphic transformation). While affine transformation alone is limited, it provides a benchmark for evaluating our full pipeline. We then calculated pairwise distances for each of the four skeletonized tracts (GABA and inotocin in both left and right hemispheres). Mean pairwise skeleton distances were significantly smaller following the combined affine and diffeomorphic transformation compared to only the affine transformation (**Figure 3F-G**). Specifically, mean distances were 5.40 ± 2.66 µm (s.d.) for inotocin-labeled tracts and 3.42 ± 1.73 µm for GABA-labeled tracts. This is similar to the 3.97 ± 3.65 µm distance between tracts reported for the most recent *Drosophila melanogaster* reference brain^10^, indicating that our clonal raider ant reference brain and registration pipeline perform at a comparable level.

### Registrations to the reference brain facilitate comparisons across experiments

To illustrate the utility of the reference brain in conducting comparisons across different labeling experiments, we focused on the central body of the CX. This region is notable for its dense GABAergic, serotonergic, and dopaminergic innervation (**Figure 4A**). Alignment in the common reference space revealed the laminar organization of the CBU^25,26^. The posterior CBU (**Figure 4A-F**) displayed prominent dorsal 5-HT signal (**Figure 4E**), while a narrow ventral band showed primarily GABA and TH labeling (**Figure 4C-D**). In the anterior CBU (**Figure 4G-L**), the banding pattern was inverted, with dorsal TH (**Figure 4I**) and ventral 5-HT signal (**Figure 4K**). Our results indicate that the *O. biroi* CBU is organized in multiple spatially overlapping layers with distinct innervation patterns, reminiscent of other insects^25,27^. The CBL exhibited a combination of GABAergic and serotonergic innervation throughout (**Figure 4D-E, J-K**). Similar to other ants^28^, but unlike bees ^29,30^ and flies^31^, we observed little TH input to the CBL. These results demonstrate sub-neuropil accuracy of our registration pipeline and underscore the value of the reference brain for visualizing neuroanatomy across experiments.

**Figure 4.**
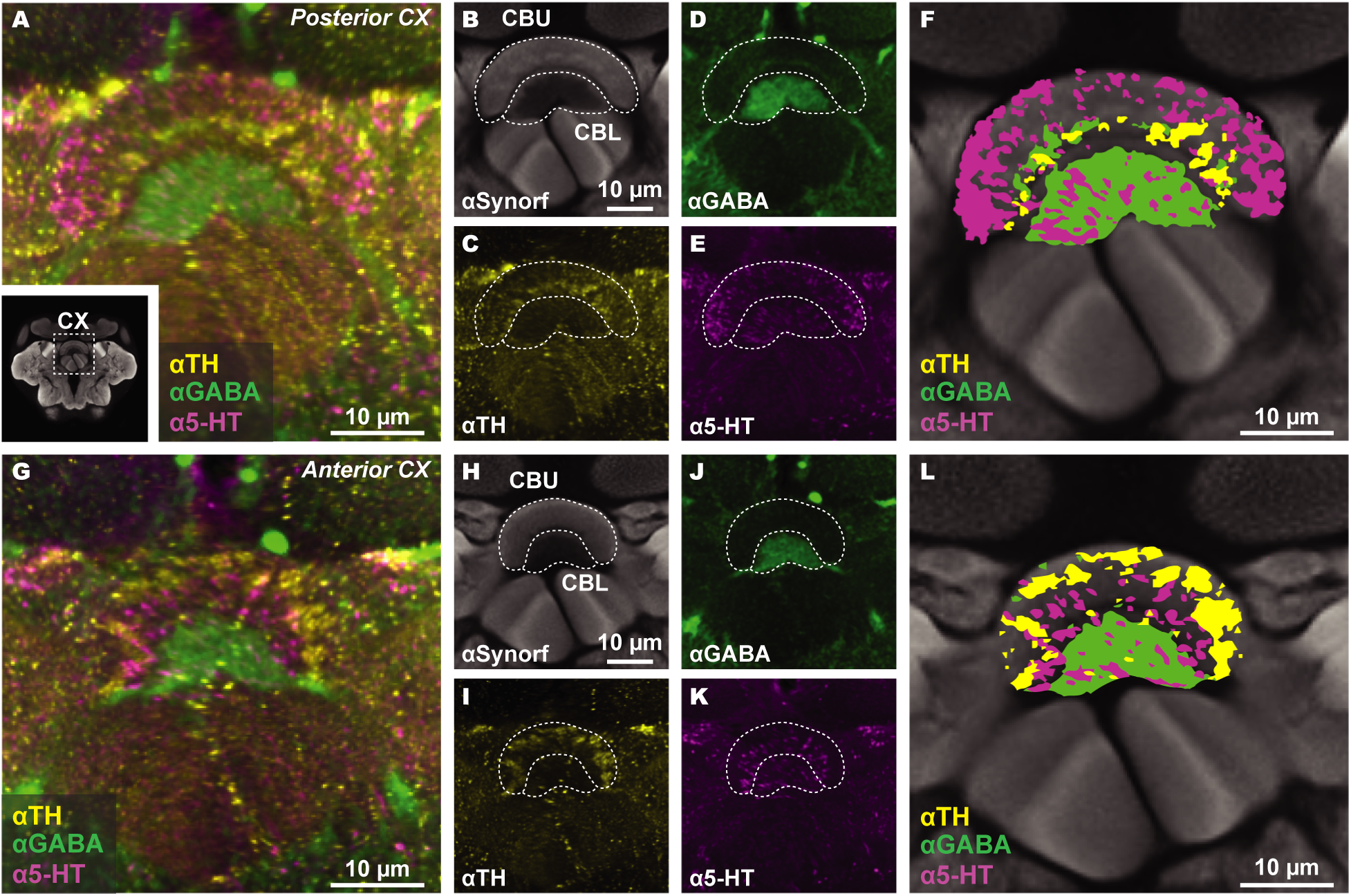
Registrations to the reference brain facilitate comparisons across experiments. (A) Composite image of the posterior CX with anti-TH, anti-GABA, and anti-5-HT staining from separate experimental brains registered to the reference brain. Inset shows the location of the CX in the brain. (B) Reference brain anti-Synapsin channel of the CX with CBU and CBL outlined (dashed lines). (C-E) Neurotransmitter channels from experimental brains displayed individually with overlaid CBU and CBL outlines from (B). (F) Camera lucida-style illustration showing CX layers and innervation pattern. (G-L) Same as (A-F), but showing an anterior position within the CX. Abbreviations: tyrosine hydroxylase (TH), gamma-aminobutyric acid (GABA), 5-hydroxytryptamine (5-HT), central complex (CX), central body upper (CBU), central body lower (CBL).

## DISCUSSION

Eusocial insects are powerful models for studying the neurobiology of complex social behavior. Here, we provide an unbiased reference brain for the clonal raider ant and a robust registration pipeline for precise comparisons across samples. This resource offers both a practical tool and a foundation for exploring the neural basis of sociality in this species.

We generated the clonal raider ant reference brain using 3D groupwise registration of 40 individual brains, an approach adapted from the JRC2018 unbiased *D. melanogaster* brain and ventral nerve cord templates^10^. This method accommodates interindividual variability and outperforms single-individual templates in registration precision and accuracy^19^. Our registration pipeline and associated GUI enable the alignment of any brain labeled with the anti-Synapsin antibody to the reference with precision comparable to the *D. melanogaster* brain template, supporting a similarly broad range of applications. Although we used immunohistochemistry as a proof-of-concept, the registration workflow is compatible with other labeling methods, including *in situ* hybridization, dye fills, and genetically encoded reporters. The reference brain also offers a framework for integrating multimodal datasets, such as registering whole-brain EM volumes to light-microscopy atlases^11,16^.

Generating a reference using 40 individual brains provided insights into interindividual variation in *O. biroi*. We observed a left–right asymmetry in the tilt of the mushroom body medial lobe, a phenomenon not previously reported in ants or other insects, but reminiscent of the *D. melanogaster* asymmetric body^32,33^. We also discovered variation in total brain volume among individuals. In other ant species, differences in brain morphology are linked with worker caste or task specialization ^34–37^, suggesting that our findings may reflect behavioral individuality and division of labor in the clonal raider ant. Future work can explore this putative connection, as well as how variation in brain architecture relates to established correlates and regulators of division of labor in *O. biroi*, including pheromone perception^38^ and neuromodulator signaling^39^.

We encourage researchers investigating the clonal raider ant to integrate our reference brain, anti-Synapsin staining protocol, and registration pipeline for neuroanatomical and gene expression analyses. By adopting a common framework, datasets collected over time can be seamlessly aligned into a shared virtual space. We envision the clonal raider ant reference brain as a foundation for generating community-driven anatomical and functional atlases, databases, and computational tools.

## ACKNOWLEDGEMENTS

We thank Stephany Valdés-Rodríguez, Alejandra Hurtado-Giraldo, Alek Rahman, and Leonora Olivos-Cisneros for ant husbandry and Yohann Chemtob for computing assistance. John Bogovic, Stephan Saalfeld, Nick Tustison and Philip Cook provided computational advice. We thank Pavel Osten and Rodrigo Muñoz Castañeda for early discussions on the reference brain and Hannah Haberkern for discussions on the CX immunohistochemistry. The anti-Synapsin antibody was developed by Erich Buchner and obtained from the Developmental Studies Hybridoma Bank, created by the National Institute of Child Health and Human Development of the National Institutes of Health and maintained at the University of Iowa, Department of Biology. Confocal microscopy was performed in part at the Rockefeller University Bio-Imaging Resource Center, RRID:SCR_017791. Some computation was performed using the Rockefeller University High Performance Computing Resource Center, RRID:SCR_025889. This work was supported by the National Institute on Deafness and Other Communication Disorders under award number K99DC021506 to D.D.F. The content is solely the responsibility of the authors and does not necessarily represent the official views of the National Institutes of Health. This work was also supported by the Howard Hughes Medical Institute via the Investigator program (to D.J.C.K.) and the James H. Gilliam Fellowships for Advanced Study program (to L.E.L. and D.J.C.K.), as well as an NSF Graduate Research Fellowship under award number DGE 194642 to L.E.L. This is Clonal Raider Ant Project paper number 39.

## AUTHOR CONTRIBUTIONS

D.D.F., L.E.L., and D.J.C.K. conceptualized and designed the study; D.D.F. and L.E.L. performed brain dissections, immunohistochemistry, and confocal imaging to generate the reference brain; L.E.L. performed 3D segmentation in Amira, completed the neurotransmitter antibody staining study, and conducted all statistical analyses; R.M. designed the reference generation and registration pipeline and wrote the associated scripts with assistance from D.D.F.; I.S. conducted the study of body-, head-, and brain size with assistance from L.E.L.; I.L. skeletonized neurite tracts with support from L.E.L. and assisted D.D.F. in brain tilt measurements; D.D.F. and L.E.L. prepared the figures with input from D.J.C.K.; D.D.F. drafted the manuscript with help from L.E.L.; D.J.C.K. supervised the project. All authors discussed the results, edited the manuscript and approved the final version.

## DECLARATION OF INTERESTS

The authors declare no competing interests.

---

**Figure S1.**
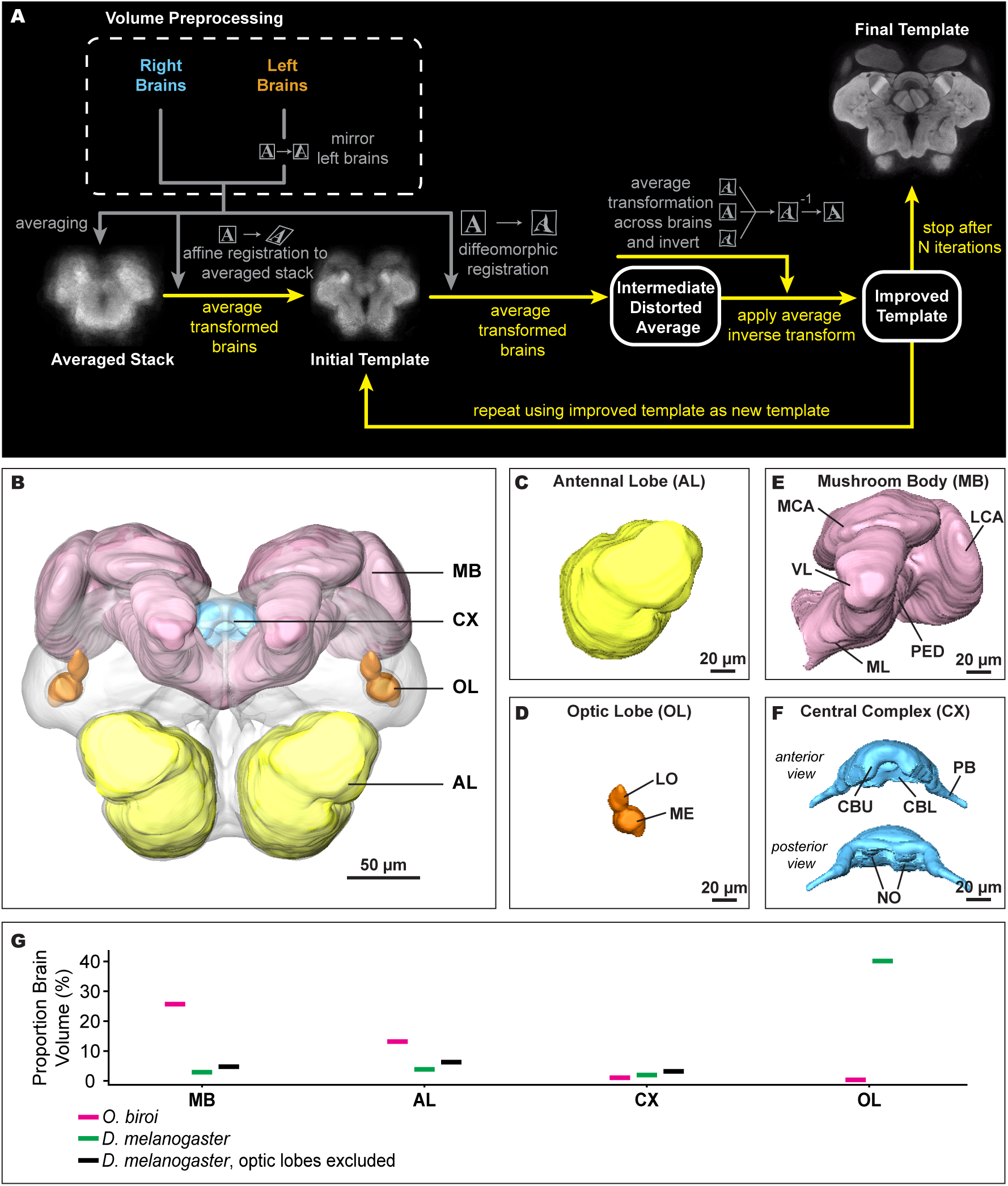
Process flowchart and anatomy of the reference brain. (A) Overview of the confocal volume preprocessing and template generation. Additional details, original code, and a tutorial to reproduce results are available on our GitHub. (B) Volume segmentation of focal neuropils using the clonal raider ant reference brain. Neuropil regions were segmented manually using the Amira 3D Pro segmentation editor tool. (C-F) Individual focal neuropils in (B) shown in greater detail. The central complex in (F) is displayed twice, once from anterior and once from posterior perspective. (G) Proportion of total brain volume occupied by the MB, AL, CX, and OL in D. melanogaster and O. biroi. For D. melanogaster, proportions are calculated using the total brain volume with and without the OLs. Abbreviations: antennal lobe, AL; mushroom body, MB; optic lobe, OL; central complex, CX; medial calyx, MCA; lateral calyx, LCA; pedunculus, PED; vertical lobe, VL; medial lobe, ML; lobula, LO; medulla, ME; central body upper, CBU; central body lower, CBL; protocerebral bridge, PB; noduli, NO.

**Figure S2.**
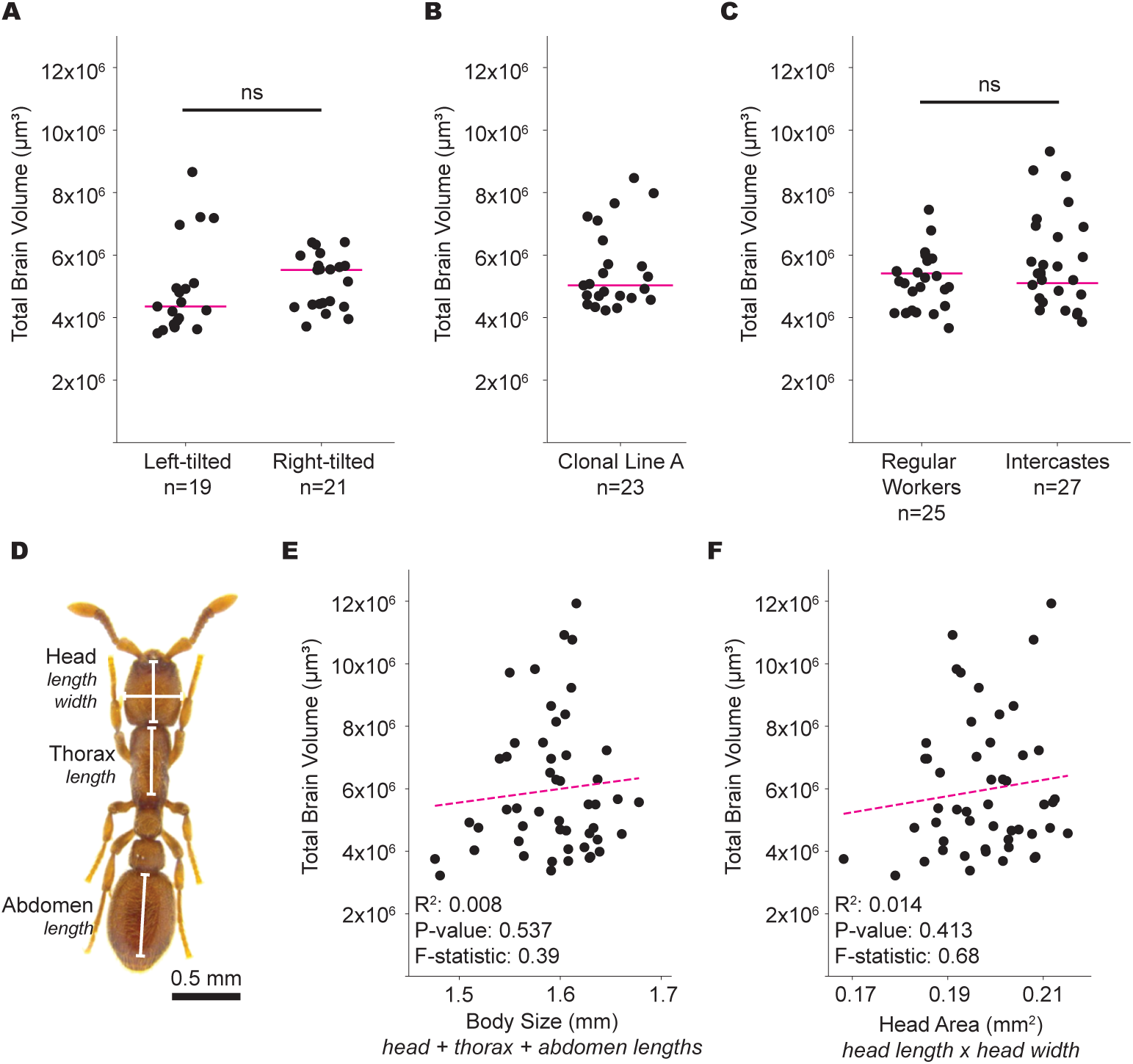
Brain volume measurements across phenotypic and genotypic groups. (A) Total brain volumes of the 40 individual brains used to generate the reference, sorted by direction of mushroom body tilt. Each data point represents a single brain; the magenta line indicates the median. Medians were compared with the Mann-Whitney U test (p = 0.144) and variances were compared using the Brown-Forsythe test (p = 0.249). (B) Total brain volumes of 23 brains from clonal line A ants. Each data point represents a single brain; the magenta line indicates the median. (C) Total brain volumes of 52 brains from clonal line B ants, sorted by whether the individual was a regular worker or an intercaste. Each data point represents a single brain; the magenta line indicates the median. Medians were compared with the Mann-Whitney U test (p = 0.164) and variances were compared using the Brown-Forsythe test (p = 0.08). (D-F) Analysis of total brain volume as it relates to ant body size and head area for 50 clonal line B ants. (D) Photograph of a clonal raider ant worker illustrating the morphometric measurements used to quantify body size and head area. (E) Linear regression plot of total brain volume (Y axis) and the estimated body size of ants (X axis). The body size estimate is equal to the sum of the head, thorax, and gaster lengths. Each data point represents a single brain; the magenta line indicates the median. (F) Linear regression plot of total brain volume (Y axis) and the estimated head area of ants (X axis). The head area estimate is equal to the product of the head width and head length. Each data point represents a single brain; the magenta line indicates the median.

## MATERIALS AND METHODS

### Experimental model details

Stock colonies of the clonal raider ant, *Ooceraea biroi*, were maintained in constant light at 25°C in Tupperware containers (40 x 26 cm) with a ∼2 cm thick plaster of Paris floor. During the brood care phase, colonies were fed 3 times weekly with frozen fire ant (*Solenopsis invicta*) brood and cleaned and watered once per week, as necessary. Ants used in our immunohistochemistry experiments were removed from stock colonies as newly eclosed callows, transferred to smaller (14 x 14 cm) plastic containers with a moist plaster of Paris floor, and fed and watered as described above. These conditions were maintained for 1 month, after which ants were dissected. For the intercaste vs. regular worker comparison, we generated a higher proportion of intercaste individuals by increasing our feeding schedule to 5 times per week. Newly eclosed callows were then separated and aged for 1 month following the protocol above. Before brain dissections, intercastes were identified based on the presence of vestigial eyespots^40,41^.

### Sample preparation and immunohistochemistry

Ants were anesthetized on ice before dissecting. Brains were dissected in cold 1X PBS using sharp forceps and transferred to a 1.5mL tube containing ∼200µL of 1X PBS with 4% PFA fixative on ice. After one hour of dissections, the solution was replaced with fresh fixative, and the tubes were transferred to a nutator mixer, where the brains were fixed for 2 hours at room temperature. After fixation, brains were washed three times in fresh 1X PBS and stored at 4°C.

All antibody staining performed in this study followed the same protocol. Brains were blocked for 1 hour in 1X PBS containing 5% normal donkey serum and 0.05% Triton-X 100 (hereafter referred to as blocking buffer). After blocking, brains were transferred to fresh blocking buffer containing the appropriate primary antibodies (see Supplemental Table 1) and allowed to incubate for 4 days at room temperature on a nutator mixer. After primary incubation, brains were washed three times in 1X PBS and incubated in fresh blocking buffer containing secondary antibodies conjugated to Alexa Fluor probes and DAPI counterstain. Secondary incubation proceeded for 3 days, after which brains were again washed three times in fresh PBS and stored at 4°C. Brains were mounted on silane-coated slides in Slowfade Glass mounting media.

### Confocal imaging

Confocal microscopy of antibody-stained brains was conducted using Zen image acquisition software on a Zeiss LSM 880 and a Zeiss LSM 900 equipped with 405nm, 488nm, 561nm, and 633nm laser lines. The 40 brains in our reference brain dataset were imaged on the Zeiss LSM 880 with AiryScan using a Zeiss LD LCI Plan-Apochromat 40X / 1.2NA objective and Zeiss Immersol-G immersion media. Image tiles were acquired at 0.1289 x 0.1289 x 0.295 μm^3^ voxel resolution and processed and stitched using Zen Blue image processing software. All other samples were imaged on a Zeiss LSM 900 using a Zeiss LD LCI Plan-Apochromat 40X / 1.2NA or a Zeiss Plan-Apochromat 20X / 0.8 NA objective.

### Reference brain construction

Unless otherwise noted, all confocal volumes were processed with Advanced Normalization Tools (ANTs) library^20,21^ using custom Python scripts. First, all 40 individual confocal volumes were opened in FIJI/ImageJ and aligned in the same dorsal-ventral and anterior-posterior orientation, and extra space was added in each dimension to prevent clipping. All brains were then designated as having a left or a right tilted mushroom body based on visual inspection of the angle where the medial lobes of the two hemispheres meet at the midline. In addition to the angle where the medial lobes meet, we also noted a difference in the staining intensity and relative position of a small band of neuropil whose orientation appeared to demarcate the direction of the tilt. However, the identity of this region within the mushroom body is not obvious and we did not investigate this further. All left-tilted brains were mirrored in ANTs to produce a dataset composed solely of right-tilted brains. Next, all volumes were resampled to 0.4 μm^3^ isotropic voxels using ANTs. Our reference was generated using the ANTs antsMultivariateTemplateConstruction.sh script. First, we used a single iteration of affine registration (maximizing mutual information) onto the unregistered average of all pre-processed volumes and averaged all the registered brains to generate the initial affine template. We then used a cross-correlation-maximizing diffeomorphic GreedySyn registration-based template building pipeline^22^ using a progressive multiscale approach with 60, 180, 40 and 20 iterations at 1/8,1/4,1/2, and original scale respectively. We also generated a reference brain with a voxel resolution of 0.8 μm^3^ via resampling, which was used to segment neuropils due to its smaller file size. All reference brain construction code and a detailed tutorial for reproducing our results can be found on the GitHub repository^42^.

### Segmentation and analysis of 3D neuropil structures

The reference brain and individual brains were segmented using Amira 3D Pro software in the anti-Synapsin channel. Brain volume files were downsampled to 0.8 μm^3^ in ANTs to decrease file size for processing in Amira. We performed semi-automated total brain volume segmentation using a custom Amira recipe and the output was manually refined. Focal brain neuropils were manually segmented using the Amira 3D Pro Segmentation Editor tool. To assess whether total brain volume scaled with focal neuropil volume, we performed linear regression using ordinary least squares (OLS). Total brain volume was used as the dependent variable, and focal neuropil volume was used as the independent variable. For bilaterally symmetric structures, we used the average volume of the left and right hemispheres. Model fit was evaluated by computing the coefficient of determination (R²), slope, intercept, p-value, standard error, and F-statistic. Full regression results are reported in the corresponding figure panel and caption.

### Body and head size measurements

We conducted morphometric measurements to quantify the size of the workers. Ants were immobilized under a piece of acrylic in a standardized posture and photographed using a Leica MSV266 brightfield microscope. A ruler was photographed under identical imaging conditions for calibration. Body size was calculated by summing the length of the head, thorax, and abdomen. Head length was defined as the length of the line bisecting the head from the clypeus to the posterior-most point of the head, while head width was defined as the length of the line running medial-lateral positioned at the widest part of the head (Figure S2E). Head area was calculated by multiplying the head length by the head width. To assess whether total brain volume scaled with body size, we performed linear regression using ordinary least squares. Body length or head width was used as a continuous independent variable, and total brain volume served as the dependent variable. Model fit was evaluated by calculating the coefficient of determination (R²), slope, intercept, p-value, standard error, and F-statistic. All statistical results are reported in the respective figure panels and captions.

### Registering confocal volumes to the reference

Two-channel confocal volumes of brains labeled with an anti-Synapsin antibody and a second neurotransmitter antibody were registered to the 0.4 μm^3^ isotropic voxel reference brain. First, confocal stacks were rotated in Fiji/ImageJ so that the axes of the image to be registered aligned with that of the reference brain. Next, the stacks were resampled in ANTs to match the 0.4 μm^3^ isotropic voxels of the reference brain. A Contrast Limited Adaptive Histogram Equalization (CLAHE) was then applied to the anti-Synapsin channel in Fiji. We generated user-friendly registration and warping GUIs to run ANTs scripts that register brains to the reference. Using the registration GUI, the anti-Synapsin channel was first registered to the reference brain using an affine transformation. The GUI includes a setting for the left-tilted brains, where the brain is mirrored before registration. The affine-transformed anti-Synapsin volume was then registered to the reference brain using a diffeomorphic GreedySyn transformation and a cross-correlation similarity metric, also using the registration GUI. We empirically determined that iterations of 30, 30, 30, 90, 20, and 8 at 1/32, 1/16, 1/8, 1/4, 1/2, and original scale, respectively, worked sufficiently for registration of most brains. Following registration, both the affine and diffeomorphic transformations were applied to the neurotransmitter channel using the warping GUI. Both anti-Synapsin and neurotransmitter channels were visualized in Fiji. All ANTs scripts, the GUIs for registering samples to our reference brain, and a tutorial can be found on GitHub^42^.

### Skeletonization of GABA and inotocin neurite tracts

We identified conspicuous neurite tracts visible via antibody staining for GABA and inotocin that were suitable candidates for skeletonization. For each of the brains stained for GABA (n=8) and inotocin (n=6), we manually skeletonized the left and right neurite tract using the Simple Neurite Tracer (SNT)^43^ Fiji toolbox. Skeletons were generated for the affine-transformed and diffeomorphic-transformed neurotransmitter channels. To quantify anatomical similarity between skeletons across brains, we calculated pairwise distances for each tract and transformation type. For each pair of skeletons, we extracted the 3D coordinates from SWC files and computed a bidirectional average minimum distance. Specifically, for two skeletons A and B, we calculated the full pairwise Euclidean distance matrix between all points in A and B. We then determined the mean of the minimum distances from each point in A to its nearest neighbor in B, and vice versa. The final pairwise distance was defined as the average of these two directional means, providing a symmetric metric that captures both spatial deviations and local mismatches in morphology. All pairwise distances within each group (e.g., GABA-left-affine) were calculated, and mean pairwise distances were compared across registration conditions using Welch’s two-sample *t*-test to account for unequal variances.

### Quantification and statistical analysis

All statistical analyses were performed using custom scripts in Python/Jupyter Notebook. The details for statistical tests, including the type of test used, the exact sample size n, the definition of n, the definition of center, and dispersion and precision measures can be found in the figures and figure legends. Statistical significance was defined as a p-value < 0.05.

Our GitHub repository^42^ contains an in-depth tutorial for reproducing our results, as well as a user-friendly GUI and detailed instructions for registering additional samples to the clonal raider ant reference brain. Original Python scripts used for statistical analyses and plotting are available in a dedicated GitHub repository ^44^ and will also be made available as a Zenodo repository immediately upon publication.

---

**Table S1.** Immunohistochemistry antibodies and description of anatomical labeling. Abbreviations: DSHB, Developmental Studies Hybridoma Bank; AL, antennal lobe; MB, mushroom body; LH, lateral horn; CX, central complex; OL, optic lobes; SEZ, suboesophageal ganglion; VNC, ventral nerve cord.

| Antigen | Antibody identifier | Vendor | Host | Results in *O. biroi* brain | Used in this study? |
| --- | --- | --- | --- | --- | --- |
| Synapsin | 3C11 | DSHB | Mouse | Strongly labels all brain neuropil. No signal in axon tracts, but weak and nonspecific signal in cell body layer occasionally observed. | Yes |
| Tyrosine hydroxylase | NB300-109 | Novus | Rabbit | Labels a cell cluster dorsal to the AL and another near the MB calyx region. Smaller soma groups are associated with the CX and OLs. Widespread punctate signal is visible throughout the brain. | Yes |
| GABA | A2052 | Sigma | Rabbit | Broadly labels somas around the perimeter of the AL. Additional cell clusters associated with the LH region, calyces, and CX. Several minor axon tracts are visible. | Yes |
| Serotonin | S5545 | Sigma | Rabbit | Labels approximately 20 somas near the posterior surface and a pair of somas dorsal to the AL. Punctate signal is visible throughout most neuropils. | Yes |
| Inotocin | Custom | YenZym | Goat | Labels a pair of neurons at the posterior surface of the SEZ which predominantly innervate the perimeter of the region. Fibers appear to descend to the VNC. | Yes |
| Elav | 9F8A9 | DSHB | Mouse | No signal. | No |
| Elav | 7E8A10 | DSHB | Rat | No signal. | No |
| Repo | 8D12 | DSHB | Mouse | Nonspecific labeling of all brain somata. | No |
| Bruchpilot | NC82 | DSHB | Mouse | No signal. | No |
| Alpha tubulin | 4A1 | DSHB | Mouse | Labels axonal tracts, but with signal also visible in neuropil and cell body layer. Antennal nerve and sensory innervation of the antennal lobe is visible. Poor label in deeper brain regions. Preferable over 12G10. | No |
| Alpha tubulin | 12G10 | DSHB | Mouse | Weakly labels tracts, but with signal also visible in neuropil and cell bodies. Poor label in deeper brain regions. | No |
| Acetylated alpha tubulin | T7451 | Sigma | Mouse | Labels axonal tracts, but with signal also visible in neuropil and cell body layer. Antennal nerve and sensory innervation of the antennal lobe is visible. Poor label in deeper brain regions. | No |
| Beta tubulin | E7 | DSHB | Mouse | No signal. | No |
| Serotonin | 20080 | Immunostar | Rabbit | No signal. | No |
| Choline acetyl transferase | ChAT4B1 | DSHB | Mouse | Labeling throughout the brain. | No |
| Histamine | 22939 | Immunostar | Rabbit | No signal. | No |

## Notes

### Competing Interest Statement

The authors have declared no competing interest.

